# Intra-ocular dendritic cells are increased in HLA-B27 associated acute anterior uveitis

**DOI:** 10.1101/2021.02.16.431370

**Authors:** Maren Kasper, Michael Heming, David Schafflick, Xiaolin Li, Tobias Lautwein, Melissa Meyer zu Horste, Dirk Bauer, Karoline Walscheid, Heinz Wiendl, Karin Loser, Arnd Heiligenhaus, Gerd Meyer zu Horste

**Affiliations:** Ophtha-Lab, Department of Ophthalmology, and Uveitis Centre at St. Franziskus Hospital, Muenster, Germany; Department of Neurology with Institute of Translational Neurology, University Hospital Muenster, Muenster, Germany; Augen-Zentrum-Nordwest Augenpraxis Ahaus, Ahaus, Germany; Department of Ophthalmology, University of Duisburg-Essen, Essen, Germany; Department of Human Medicine, University of Oldenburg, Germany; University of Duisburg-Essen, Essen, Germany

**Keywords:** HLA-B27, Uveitis, Inflammation, Dendritic Cells

## Abstract

Uveitis describes a heterogeneous group of inflammatory eye diseases characterized by infiltration of leukocytes into the uveal tissues. HLA-B27-associated uveitis is a common subtype of uveitis and a prototypical ocular autoimmune disease. Local immune mechanisms driving human uveitis are poorly characterized mainly due to the limited available biomaterial and subsequent technical limitations.

Here, we provide the first high-resolution characterization of intraocular leukocytes in HLA-B27 positive and negative anterior uveitis and an infectious endophthalmitis control by combining single cell RNA-sequencing (scRNA-seq) with flow cytometry and low-input proteomics. Ocular cell infiltrates consisted primarily of lymphocytes in uveitis and of myeloid cells in infectious endophthalmitis. HLA-B27 positive uveitis exclusively featured an plasmacytoid and classical dendritic cells (cDC) infiltrate, and of plasma cells. Moreover, cDCs were central in predicted local cell-cell communication.

The data suggests a unique pattern of ocular leukocyte infiltration in HLA-B27 positive uveitis with relevance of particular dendritic cells.

## Introduction

Uveitis describes a heterogeneous group of inflammatory diseases involving the intraocular cavity of the eye that is devoid of immune cells under non-diseased conditions. Non-infectious uveitis is supposed to be an autoimmune disorder probably associated with various immune-mediated systemic diseases. According to the Standardization of Uveitis Nomenclature (SUN) working-group, uveitis is classified based on the anatomical location of uveitis as anterior, posterior, intermediate, and panuveitis^1^. Anterior uveitis (AU) with primary site of inflammation in the iris and ciliary body is its most frequent type (approximately 50% of cases) probably resulting in permanent vision loss through e.g., secondary cataract, glaucoma or macular edema^2^.

Acute anterior uveitis (AAU) associated with the HLA haplotype B27 (B27+AAU) is the most common and often severe form of uveitis. AAU typical clinical features include acute onset of discomfort, eye redness, tearing, visual impairment, and excessive cellular infiltration in the anterior chamber. The prevalence of the HLA-B27 haplotype is approximately 8-10% in the Caucasian population, and 40-70% among AAU patients^3^ and thus represents a strong genetic risk factor^4,5^. The HLA-B27 allele also conveys an increased genetic risk for other autoimmune diseases, including spondyloarthropathies (SpA) and inflammatory bowel disease (IBD). B27+AAU is thus regarded as a prototypic ocular autoimmune disease and, accordingly, its treatment is based on corticosteroids^6,7^ and classical and biological disease modifying anti-rheumatic drugs^8,9^. Notably, although T- cell-inhibition (e.g., by cyclosporine A) is beneficial^10^, it provides limited efficacy in B27+AAU^11^, suggesting a yet poorly defined role of innate immune cells in its pathogenesis.

Immune cells infiltrating the anterior chamber of the eye can be observed clinically with the use of a slit-lamp, but are difficult to *obtain* for further analyses. In fact, the aqueous humor (AqH) invasive sampling is rarely clinically justified, and is significantly altered by given anti-inflammatory treatment. Our knowledge of aqueous humor (AqH) -infiltrating leukocytes and underlying mechanisms of uveitis thus remains superficial despite AAU diseases frequency and severity. AqH fine needle aspirates currently are applied under clinical circumstances primarily for the verification of e.g. infectious uveitis, and rarely to unraveling the pathogenesis of non-infectious uveitis ^13–16^.

In AAU, a role of bacterial triggers has been proposed^17,18^. Therefore, innate immune cells such as monocytes and dendritic cells (DC) that phagocytose and process foreign antigens are of special interest. In fact, phenotypic analysis of circulating monocytes in patients with different subtypes of autoimmune-mediated uveitis emphasized phenotypic differences during the disease course^19–22^. The microbiome, and other potential mechanisms accounting for dysbiosis, barrier dysfunction and immune response have also been suggested to contribute to the pathogenesis of AAU^23^. Soluble mediators^24–27^ and infiltrating leukocytes have been analyzed^28^ previously in the AqH of uveitis patients, but access to samples and technical challenges remain the main bottlenecks towards better understanding pathomechanisms in uveitis.

Here, we combined single cell RNA-sequencing (scRNA-seq) with confirmatory flow-cytometry and low input proteomics of soluble cytokines and thereby provide the first unbiased characterization of leukocytes in the AqH from patients with B27+AAU compared to patients with HLA-B27 negative AU (B27-AU), and acute infectious endophthalmitis. We found that lymphocytes predominate in the intraocular infiltrate in B27+AAU. Notably, B27+AAU showed an elevated frequency of plasmacytoid and classical DC. In B27+AAU, DC has the highest amounts of predicted intercellular interactions and increased expression of AAU- and SPA-related GWAS risk genes that distinguished this uveitis from active B27-AU, suggesting a specific involvement of these cell types in B27+AAU. Subtypes of anterior uveitis thus show specific patterns of local leukocyte responses.

## Results

### Patients recruitment, samples collection, and optimizing analysis of aqueous humor (AqH)

We here aimed for an unbiased characterization of leukocytes in the AqH in anterior uveitis flares using sc transcriptomics and confirmatory techniques. We included eleven patients with current onset of uveitis flare (n=8), or endophthalmitis (n=3) into this study, being untreated with topical corticosteroids for this flare (Suppl. Tab.1). Out of 4,980 total patients with any intraocular inflammation seen at our uveitis center in that period; corresponding to a recruitment rate of approximately 0.18% (Suppl. Tab.1). Processing failed technically in four samples, leaving seven samples for final sc-RNA-seq analysis, including four patients with active B27+AAU, two patients with active B27-AU, and another one with active bacterial endophthalmitis *(Streptococcus pneumoniae).* In four patients with sc-RNA-seq, cells were analysed further via flow cytometry, and in 10 of the AqH samples cytokine analysis was also performed.

The endophthalmitis patients were significantly older than B27+AAU patients (ANOVA p=0.0157; Suppl. Tab. 1). There were four females in the B27+AAU cohort, and three males and one female in the B27-AU cohort. All B27+AAU patients had associated SpA. Both uveitis groups did not differ significantly regarding age (ANOVA p=0.566), ANA status, frequency of systemic anti-inflammatory treatment, topical medication, previous ocular surgery or uveitis duration.

### Single cell transcriptomics reconstructs key leukocyte lineages in aqueous humor

We then performed scRNA-seq of infiltrating cells from rare AqH fine needle aspirates (Suppl.Tab.1). Thereby, we obtained transcriptional information of 13,550 total cells, and 1,936 average cells (±1,411 SD) per patient with 830 average genes (± 402 SD) detected per cell (Suppl. Tab.2). After quality control, we clustered sc transcriptomes of all patients combined and identified 13 individual cell clusters (Fig. 1A,B). We manually annotated these clusters to cell types based on the expression of canonical marker genes (Fig. 1C, Suppl. Fig.2, Suppl. Tab.3). As previously shown in AqH aspirates of human uveitis patients^28^, only hematopoietic cells were identified among these clusters. Grossly, clusters were classified into cells of myeloid (40%; cDCa, cDCb, pDC, matDC, granulo, myeloid) and of lymphoid origin (60%; NK, gdTC, CD8, Treg, CD4, naive Bc, plasma). Myeloid clusters separated into classical DC clusters named cDCa *(ITGAX, CLEC7A)* and cDCb *(CLEC10A, MRC1),* plasmacytoid (p)DC *(IL3RA/CD123, CLEC4C/CD303)* and mature (mat)DC *(TMEM176B, IDO1, FSCN1, LAMP3, CD83),* granulocytes (granulo; S100A12 / *S100A8 high, CCL2 low)* and myeloid cells with unclear assignment (myeloid; S100A12 / *S100A8 low, CCL2 high)* (Fig. 1C). Lymphoid clusters were classified as CD4+ T-cells (CD4; *IL7R, CD3G),* CD8+ T-cells (CD8; *CD8A, CD3G),* γδ T-cells (gdTC; *NKG7, TRDC),* regulatory T-cells (Treg; *IL2RA, FOXP3),* natural killer cells (NK; *GNLY, NKG7),* naive B-cells (naive BC; *MS4A1, CD19, IGHD)* and plasma cells (plasma; *IGHG1, CD38, SDC1/CD138)* (Fig. 1C). We thus successfully identified all major leukocyte lineages in AqH fine needle aspirates. The relative cluster composition (Fig. 1B, Suppl. Tab.4) was considerably different from peripheral blood described previously^29^.

**Figure 1:**
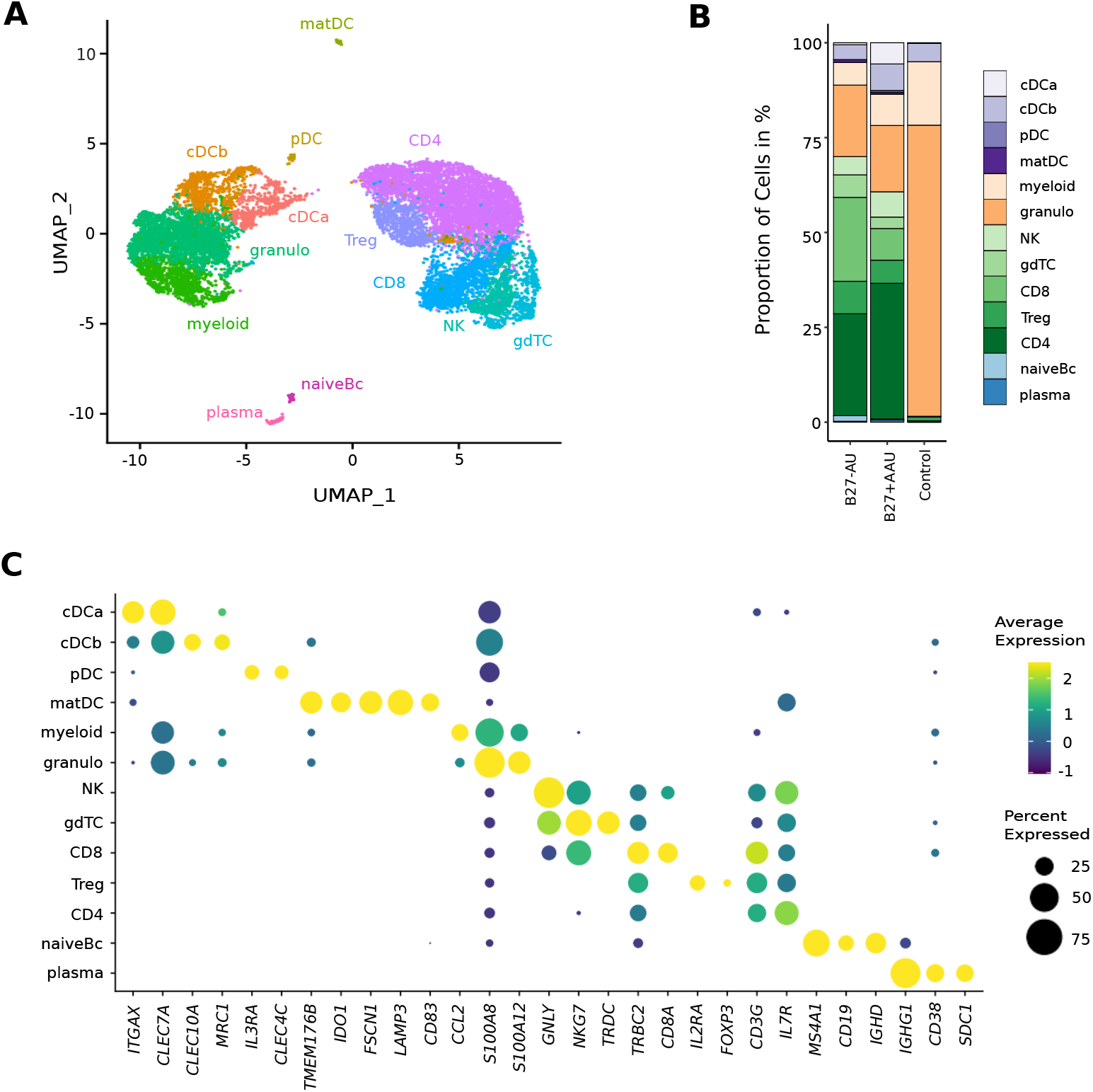
Single cell transcriptomics reconstructs leukocyte subsets infiltrating the eye. (A) UMAP projection of pooled 7 samples (control n=1; B27-AU n=2; B27+AAU n=4). The sc transcriptomes were manually annotated to cell types based on marker gene expression, and distinguished in 13 cell clusters (color-coded; each dot represents one cell). (B) The mean proportion of cells (%) in each cluster per group is depicted in a stacked bar-plot. (C) Dot plot of selected marker genes grouped by cluster. The average gene expression level is color-coded and the circle-size represents the percentage of cells expressing the gene. Threshold was set to a minimum of 10% of cells in the cluster expressing the gene. DC: dendritic cell, pDC: plasmacytoid cell, matDC: mature DC; granulo: granulocytes, NK: natural killer cells, gdTC: γδ T-cells, Treg: regulatory T-cells, Bc: B-cells.

### Diverse uveitis entities exhibit a unique intraocular leukocyte composition

We next sought to understand how intraocular inflammation differed between uveitis entities. First, we assessed the endophthalmitis control patient and found almost exclusively myeloid lineage clusters with predominating granulocytes in accordance with an acute anti-bacterial response (Fig. 2B; Suppl. Fig. 1)^30^.

**Figure 2:**
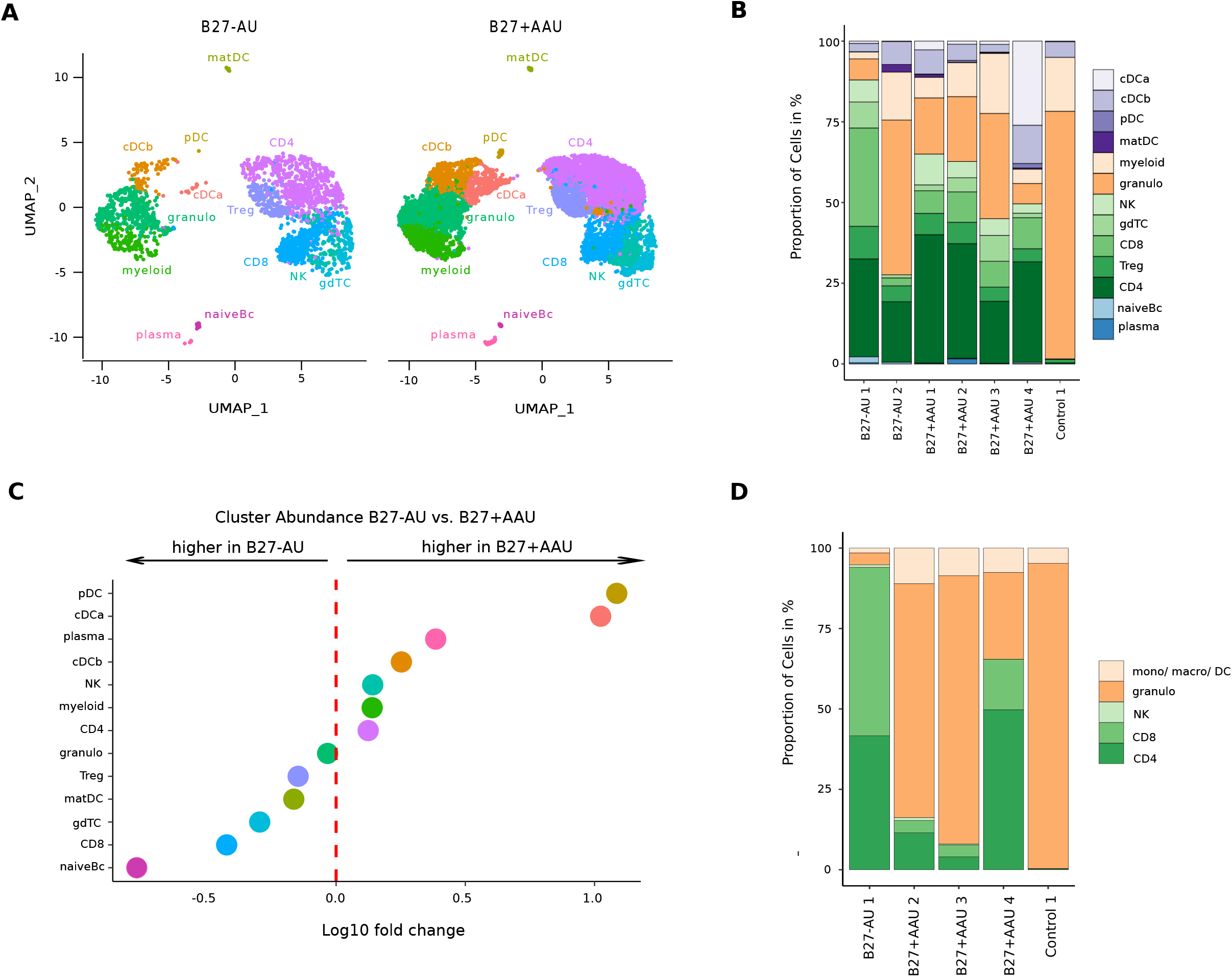
A unique intraocular leukocyte pattern characterizes individual uveitis causes. (A) UMAP projection of pooled B27-AU (n=2) versus pooled B27+AAU (n=4) samples. The sc transcriptomes were manually annotated to cell types based on marker gene expression, and distinguished in 13 cell clusters (color-coded; each dot represents one cell). (B) The proportion of cells (%) in each cluster is depicted in a stacked bar-plot for individual samples. (C) Dot plot of cluster abundance of B27-AU versus B27+AAU. X axis represents the decadic logarithm of fold change of proportional cluster size. (D) Leukocytes of AqH samples were analyzed according to their frequency [%] of granulocytes, monocytes / macrophages / DC, CD4+ and CD8+ T-cells, and NK-cells. The proportion of each cell population identified via flow cytometry is depicted in a stacked bar plot. DC: dendritic cell, pDC: plasmacytoid cell, matDC: mature DC; granulo: granulocytes, NK: natural killer cells, gdTC: γδ T-cells, Treg: regulatory T-cells, Bc: B-cells, mono: monocyte, macro: macrophage.

In contrast, the cellular infiltrate in B27-AU and B27+AAU patients’ AqH was of considerably more lymphoid origin reflecting their autoimmune etiology (Fig. 2A,B). When systematically comparing B27+AAU vs. B27-AU samples, inter-patient variability was high (Fig. 2A,B), but several populations still differed between uveitis subtypes (Fig. 2C, Suppl. Tab.4). While the CD8 and naiveBc clusters were reduced, the pDC and cDCa clusters were considerably more abundant in B27+AAU patients (Fig. 2C) overall indicating an influx or expansion of DC in this uveitis entity.

Next, we sought to confirm our findings using flow cytometry (Fig. 2D; Suppl. Fig. 3). AqH-derived cells of some of the patients (Suppl. Tab.1) were analyzed using key lineage markers to distinguish granulocytes (CD3^-^CD11b^+^HLA-DR^-^), monocytes/macrophages/DC (CD3^-^CD11b^+^HLA-DR^+^), CD4^+^ and CD8^+^ T-cells (CD3^+^CD11b^-^), and NK-cells (CD3^-^CD11b^-^CD56^+^). The analysis confirmed the mainly myeloid infiltrate in the endophthalmitis patient (Fig. 2D). B27-AU patients showed an infiltrate dominated by T-cells, low NK-cells and low cells of myeloid origin. In B27+AAU patients, myeloid cells were more abundant than in B27-AU (Fig. 2D). Also, the abundance of gross cell types quantified by scRNA-seq and flow cytometry showed high positive correlation (Fig. 2D, Suppl. Fig. 3B). The higher frequency of granulocytes detected in flow cytometry than in the scRNA-seq might be due to higher fragility of these cells during processing for scRNA-seq^31^. Overall, diverse subtypes of uveitis are characterized by a unique pattern of local inflammatory cells.

### Specific transcriptional phenotype of intraocular leukocytes in subtypes of uveitis

Next, we sought to understand how local leukocytes differ transcriptionally in diverse uveitis entities. To analytically account for low total cell numbers, we merged clusters into five broad cell type ‘metaclusters’ only for differential expression analysis (Fig. 3A, Suppl Tab.5): helper T-cells *(help;* Treg, CD4), cytotoxic cells (*toxic*; CD8, NK, gdTC), merged DC (*mergeDC*; matDC, pDC, cDCa, cDCb), other myeloid cells (*myeloidLin*; myeloid, granulo) and B-cell lineage (*BcLin*: naiveBc, plasma). We then tested for differentially expressed (DE) genes between B27+AAU vs. B27-AU. Across all clusters, multiple MHC class I and class II related genes *(HLA-A, HLA-DPA1, HLA-DRA, HLA-DRB1)* were expressed at lower level in B27+AAU samples (Fig. 3B-F). In these B27+AAU, the *help* metacluster downregulated signs of class I-dependent antigen presentation *(HLA-A, HLA-C, B2M)* and one cytokine receptor *(CXCR4).* The *toxic* meta-cluster downregulated signs of cytotoxicity *(GZMK, GZMH, LTA*). The BcLin meta-cluster featured an increase of macrophage-inhibitory factor (*MIF*), responsible for inhibition of NK-cell activity^32^, while genes related to antigen presentation (*B2M, HLA-A, HLA-DRB1)* were down regulated. The *myeloid* meta-cluster downregulated signs of class I and II-mediated antigen presentation (*HLA-A*, *HLA-DRB1),* and several interferon-regulated genes *(IFITM* genes; Fig. 3B-F).

**Figure 3:**
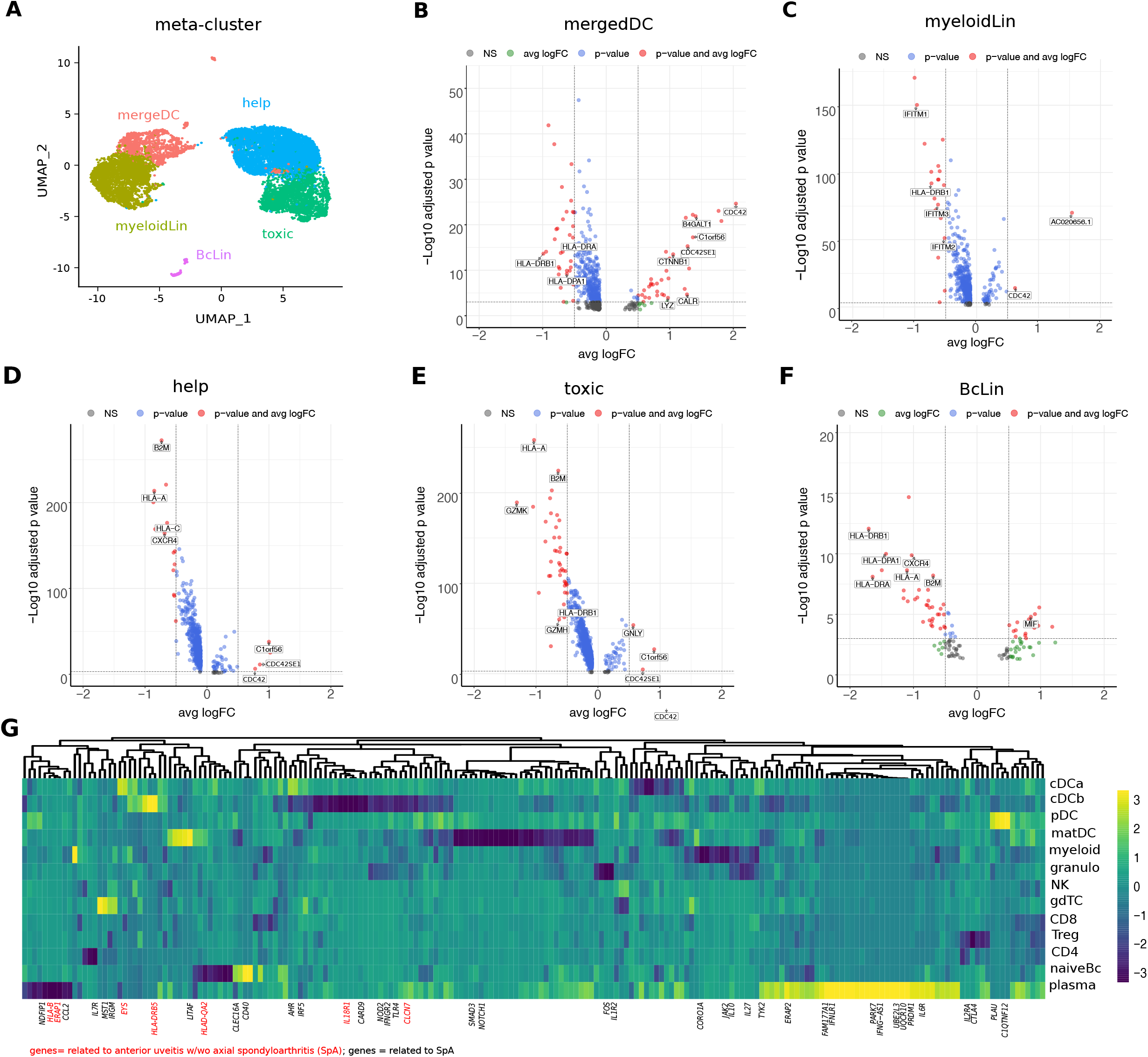
Intraocular leukocytes express a subtype-specific transcriptional phenotype. (A) UMAP projection of pooled B27-AU (n=2) versus pooled B27+AAU (n=4) samples. The sc transcriptomes were manually annotated to cell types based on marker gene expression, and distinguished in 5 metacluster (color-coded; each dot represents one cell). DE genes of B27-AU vs. B27+AAU of each meta-cluster (B) *mergeDC* (matDC, pDC, DCa, cDCb), (C) *myeloidLin* (myeloid, granulo), (D) *help* (Treg, CD4), (E) *toxic* (CD8, NK, gdTC), (F) and *BcLin* (naiveBc, plasma) are depicted as volcano plot. The threshold for average log fold change (avg logFC) was set to 0.5 and for adjusted p value to 0.001. Selected genes are labeled. (G) Heatmap showing differences in average gene expression (B27+AAU minus. B27-AU) of GWAS risk genes (Suppl. Tab. 7). Data were scaled column wise. Columns are clustered using euclidean distance measure and complete linkage. Yellow color indicated a higher expression in B27+AAU samples, blue color indicates higher expression in B27-AU samples. Genes labeled in red are anterior uveitis related, genes labeled in black SpA related risk genes.

Elevated *CTNN1* expression in mergeDC (Fig. 3B) point to an involvement of WNT catenin pathway as previously described for SpA^33^. Increased expression of *Lyz* and its cognate antisense *AC020656.1,* and *CALR^34^* and its antisense *AC092069.1* within the mergeDC cluster may suggest an activation of myeloid cells and simultaneously local counter regulating mechanisms to preserve immune privilege in the anterior chamber. Elevated expression of *B4GALT1* in mergeDC, in detail cDCa and cDCb (Supp. Tab. 6), reflect the migratory capacity of DC^35^. Notably, the mergeDC, help and toxic meta-clusters all upregulated *C1ORF56,* an oncogene previously found induced in activated lymphocytes in IBD^36^ and splenic NK-cells^37^.

We also found elevated expression of prefoldin 5 (PFDN5) in *myeloid, toxic* and *help* cluster (Suppl. Tab.5), which was recently described as a specific marker for B27+AAU associated with SpA^38^. Furthermore, as previously shown in joint biopsies of SpA patients^39,40^ and in association to autoinflammatory diseases^39,40^ elevated expression of CDC42 was detected in mergeDC, myeloid, help and of CDC42SE1 in mergedDC, myeloid, help and toxic cluster (Fig 3 B-E). We next localized the expression of known genetic risk factors for AAU with / without association to SpA and associated autoimmunity to all 13 cell clusters (Fig. 3G, Supp.Tab.7). Risk gene expression in T-cell clusters was less apparent, and many risk genes were expressed across multiple clusters without apparent enrichment in individual lineages. However, the expression of autoimmune-related gene CLEC16A^41^ and costimulatory molecule CD40 was elevated in the naiveBc cluster in B27+AAU. Many risk genes were highly expressed in the plasma cluster (e.g. *ERAP2, PARK7, IL6R, IFNLR1, IFNG-AST),* in the cDCa (e.g. *IFNGR2, AHR),* cDCb *(HLA-DRB5),* and in the pDC *(PLAU; C1QTNF12)* clusters in B27+AAU (Fig. 3G, Supp.Tab.7). Notably, in B27+AAU cDCa cells showed a reduced expression of *FOS* being involved in IL-17 signaling^42^. Compared to B27-AU, expression of genes involved in the detection of pathogens (*CARD9, NOD2, TLR4)* showed lower expression in the cDCb cluster in B27+AAU. This further supports the relevance of DC subsets in B27+AAU.

### Subtype-specific local leukocyte communication in uveitis

We also attempted to understand how uveitis controlled the local inter-cellular signaling circuitry. We therefore adapted a rodent tool (CellPhoneDB^43^) to predict cell-cell interactions between human leukocytes in autoimmune uveitis from scRNA-seq data. The pDC and plasma clusters had to be excluded because of their small size in B27-AU. In the resulting analysis, the granulo and myeloid clusters had the highest number of predicted interactions (Suppl. Fig. 4A,B). In both uveitis groups most interactions were between myeloid, granulo, DC and gdTC clusters. Myeloid lineage clusters thus express the highest capacity for cell-cell interaction.

**Figure 4:**
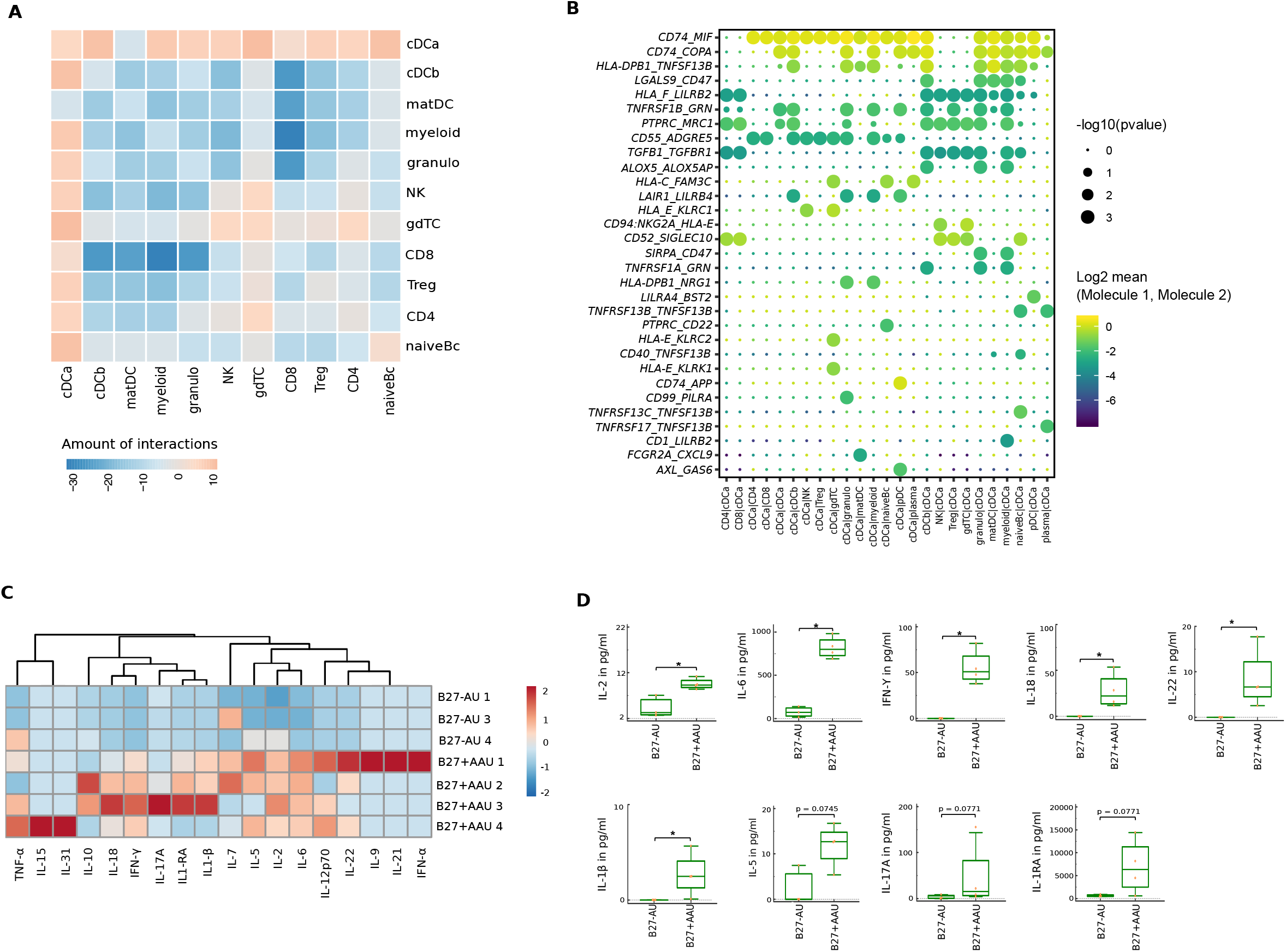
Altered local inter-cellular signalling in uveitis. (A) The total count of receptor-ligand interactions between cell clusters of B27-AU and B27+AAU sc transcriptome were obtained with CellPhoneDBv2.0 (see Suppl. Fig.4 for separate analysis). Heatmap showing the differences between B27+AAU and B27-AU (amount of predicted interactions of all cell types, excluding plasma and pDC due to few cells (<10) of B27+AAU minus those of B27-AU). (B) Overview of all ligand_receptor interactions of the cDCa cluster with at least one significant interaction. Circle size indicates the p values. The means of the average expression level of interacting molecule 1 in cluster 1 and interacting molecule 2 in cluster 2 are color-coded. (C) Heatmap showing AqH cytokine level of B27-AU and B27+AAU patients (see Suppl. Tab.8 showing single values). Data were scaled column wise. Columns are clustered using euclidean distance measure and complete linkage. (D) Boxplots of IL-2, IL-6, IFN-γ, IL-18, IL-22, IL-1β, IL-5, IL-17A, and IL-1RA (pg/ml) in the AqH of Patients with B27-AU and B27+AAU. Dots represent individual data. Mann-Whitney U-test (* p<0.05).

Next, we tested for differences of predicted interactions between both subtypes of uveitis. Therefore, we calculated the number of predicted interactions in B27+AAU minus those in B27-AU. (Fig 4A). B27-AU displayed many interactions between T/NK clusters with myeloid, matDC and cDCb clusters (Fig. 4A, Supp.Fig.2A). In contrast, the cDCa cluster showed widespread interactions in B27+AAU (Fig. 4A, Supp.Fig.4B.). This suggests that preferential interactions differ between uveitis entities and that the cDCa cluster might be centrally involved in controlling intraocular inflammation in B27+AAU.

Notably, when focussing on individual interactions of the cDCa cluster (Fig. 4B), MHC class II-related transcripts (e.g. *CD74, HLA-E)* were predicted to interact with surface molecules such as *MIF* or *KLRC1* with immunomodulatory capacity and known association to autoimmunity^44,45^. The predicted interaction of cDCa with pDC clusters in B27+AAU samples also included the *LAIR*-*LILRB4* and *AXL*-*GAS6* interaction pairs (Fig. 4B) that exert immunosuppressive phagocytosing functions^46,47^. Overall, our findings indicate subtype-specific local inter-cellular leukocyte signalling with DC forming a central signaling node in uveitis.

### Increased level of pro-inflammatory cytokines in AAU patients

Next, we sought to supplement the transcriptional data with protein information of soluble mediators. We therefore quantified a predefined set of cytokines in the AqH of these patient groups (Suppl. Tab. 8). In uveitis and endophthalmitis samples, several cytokines were below detection limits, and the most notable feature of all these samples was a pronounced presence of interleukin (IL)-6 and IL-1RA. High levels of immunosuppressive IL1-RA in AqH of uveitis/endophthalmitis patients have also been shown in endotoxin-induced uveitis (EIU) and B27+AAU patients^16,48–50^, and may reflect the immune privileged microenvironment in the anterior chamber. Corresponding with polymorphonuclear infiltrate in endophthalmitis, its cytokine milieu is characterized by innate-related IL-6, TNF-α and IL-1β (Suppl. Tab. 8) which are involved in ocular barrier breakdown and leukocyte recruitment into ocular tissue^51–53^.

Across uveitis entities, the interindividual heterogeneity of cytokine patterns was high (Fig. 4C,D). When comparing uveitis entities, the B27+AAU group showed significantly increased levels of IL-2, IL-6, IL-18, IL-22, IL-1β and IFN-γ (Fig. 4C,D). Although the AqH cytokines’ main cellular source cannot be identified in scRNA-seq, our data might reflect the activation of infiltrating myeloid cells, NK-cells and in T helper cells indicating combined Th1 (IFN-γ) and Th17 (IL-6) responses, with predominance to Th1 cells.

In addition, we then performed analysis of B27+AAU and B27-AU serum samples during the active disease stage, recruited in the context of another project^22,54^. In these samples, we found significantly elevated IL1-RA and IFN-γ levels in the serum of B27+AAU as compared to B27-AU patients (Suppl. Fig. 5, Suppl.Tab. 9) while IL-2, IL-1β, and IL-6 were absent in the serum. This indicates a major role of IL-1β, IL-6, IL-18, and IFN-γ in intraocular and IFN-γ in both intraocular and systemic immune response during uveitis activity in patients with SpA.

## Discussion

Here, we provide the first high-resolution characterization of rare and difficult to obtain intraocular leukocytes in anterior uveitis. By making scRNA-seq *per se* applicable to the AqH, we found that intraocular leukocytes are transcriptionally and compositionally different from blood, but also from other tissues affected by autoimmune diseases such as synovial fluid in rheumatoid arthritis^55^ or cerebrospinal fluid in multiple sclerosis^56^. The patient number in our study was low and inter-patient variability of observations was high. However, by comparing HLA-B27 positive versus negative anterior uveitis we identified a unique composition of intraocular leukocytes in B27+AAU. Notably, B27+AAU showed an elevated frequency of plasmacytoid and classical DC (cDC). In B27+AAU, cDC had the highest amounts of predicted intercellular interactions and increased expression of AAU- and SpA-related GWAS risk genes that distinguished this uveitis from active B27-AU. This may indicate a relevance of cDC probably promoting intraocular inflammation.

Our findings allow diverse speculations on the pathogenesis of HLA-B27 associated AAU. We find T-cells more activated in B27-AU than in B27+AAU. Indeed, SpA associated uveitis does not respond properly to T-cell inhibition^11^. Also, SpA patients (with or without AAU) mostly do not benefit from treatment with the anti-CD20 antibody rituximab^57^. In fact, AqH of AAU patients showed reduced levels of the B-cell factors BAFF and APRIL compared to patients with juvenile idiopathic arthritis-associated anterior uveitis^58^. It is notable, that naive B-cells expressing higher levels of autoimmune related CLEC16A^41^, and plasma cells, expressing several SpA risk genes, were preferentially enriched in B27+AAU. It is notable that plasma cells and the intraocular plasma cluster do not express *MS4A1/CD20.* Depletion across a wider spectrum of the B-cell lineage (e.g. by targeting CD19^59^) might be more promising. Several lines of evidence – including our study – indicate that SpA– associated B27+AAU may be driven by innate immune cells triggered through innate pattern recognition of endogenous or exogenous pathogens. Curnow et al. previously showed CD14+ monocyte/macrophage population and a separate CD14-/low BDCA-1+ (CD1c) myeloid DC population in AqH of patients with anterior uveitis^28^. Our findings lend support to a central hub function of DC.

Dendritic cells – as professional antigen-presenting cells – orchestrate the interplay between innate and adaptive immunity. The classification of DCs is complex. Traditionally, three DC subsets were defined as classical (or myeloid) DC type 1 (cDC1) or type 2 (cDC2), and plasmacytoid DC (pDC)^60^. Additional subdivisions and the definition of monocyte-derived DCs have further complexified the field (e.g.^61–63^). Most classifications are based on the expression of sets of protein markers that may or may not overlap well with transcriptional markers. In fact, a single cell transcriptomic study classified human peripheral blood DC into six subpopulations^62^: cDC1 (named DC1; *CLEC9A, C1ORF54, CADM1, CAMK2D),* CD5^high^ cDC2 cells (named (cDC2); *CD1C, FCER1A, CLEC10A, ADAMS*), CD5^low^ cDC2 cells (named DC3; *CD1D, S100A9, S100A8, VCAN, ANXA1),* monocytic-like DC (named DC4; *FCGR3A, FTL, SERPINA1, LST1, AIF1),* pDC (named DC6; *GZMB, IGJ, AK128525, SERPINF1, ITM2C),* and newly identified AXL^+^ DC DC5 *(AXL, PPP1R14A, SIGLEC6, CD22)*^64^. Notably, the phenotype and composition of tissue-resident and effector DCs is likely different from peripheral blood.

Within this pilot-study, we identified four intraocular DC clusters in two different uveitis-entities, where cDCa, cDCb and pDC share similarities to the cluster previously defined by Villani et al.^62^. We localized DC5 cells, characterized by their high expression of *AXL* and *SIGLEC6,* within the cDCa, matDC and pDC cluster. Furthermore, we identified a novel DC cluster that expressed genes related to activation and maturity (*FSCN1, CD83, LAMP3*), and was therefore designated as mature (matDC) cluster. LAMP3 was previously shown to mediate migration from tumor to regional lymph nodes and exert regulatory capacity on other cells^65^. Expression of immunregulatory *MIF* and *AXL* in cells of cDCa cluster, and their multiple predicted interactions to other infiltrating immune cells point to an immune regulatory and/or phagocytosing role of these cells during AAU. One could thus speculate that uveitis, probably in context with the microenvironment of anterior chamber-associated immune deviation includes a unique phenotype of regulatory DCs.

Pathogens are one potential trigger of autoimmune uveitis in mice, and intestinal and joint inflammation in rats overexpressing human HLA-B27 antigen, and the inflammation did not develop in a germ-free environment^66^. Transcriptional candidates identified in our study encompass a broad spectrum of functions e.g. in innate immunity, CARD9 is a candidate gene for SpA and IBD patients^67^. CARD9 has a central role in innate immunity against bacterial and fungal pathogens via the NOD2 signaling pathway. Anti-saccharomyces cerevisiae antibodies are elevated in patients with SpA and point to an involvement of CARD9 or NOD2^68^. TLR4 is a central target in EIU in rodents^69^, the related experimental model of acute anterior uveitis in humans^70^.

The S100 proteins are intracellular calcium-binding proteins and are released by cells under tissue damage, cellular stress or during inflammation, and serve as danger signals with broad spectrum effector functions^71 72 73^. S100A8/A9 and S100A12 serve as activity biomarkers of diverse autoimmune diseases^72–75^ in particular, S100A8/A9 serum levels are increased constitutively in patients with HLA-B27 associated uveitis^54,76^. In scRNA-seq, the alarmins S100A8/S100A9 are assigned mainly as marker genes for granulocytes and other phagocytes^29^. However, expression levels of S100A8/A9 did not differ between the myeloid lineage clusters of B27-AU and B27+ AAU samples.

The analysis of the cytokine pattern of AqH and sera in the study confirmed previous observations describing elevated IL-18, and IFN-γ levels in the AqH^16,77^, and elevated serum IFN-γ level in B27+ AAU patients during active uveitis^14,77^. In particular, IL-18 is a main costimulatory factor for IFN-γ expression in T- and NK-cells^78^. The cDCa cluster showed reduced *FOS* expression, mediator of IL-17 signaling^42^, pointing to a subordinated role of Th17 immunity in B27+AAU. In contrast, the cDCa cluster showed elevated expression of several IFN-γ related transcripts in B27+AAU samples, matching to previous observations in SpA joint biopsies^39^, indicating a IFN-γ driven immune response during active uveitis.

In summary, we present the first deep characterization of intraocular leukocytes by scRNA-seq in anterior uveitis patients during an ongoing flare. Our study suggests cDCs as potential central players in ophthalmic inflammation in uveitis, and it provides a basis for studying novel transcriptional candidates in greater detail in the future.

## Material and Methods

### Ethics Approval

All patients were recruited at the Department of Ophthalmology at the St. Franziskus Hospital, Muenster, Germany. The study protocol was approved by the local ethics committee of the Medical Association Westphalia-Lippe (AEKWL approval ID 2017-017-f-S). Due to the invasiveness of anterior chamber puncture, approval was limited to a total of eleven patients. The study was performed in accordance with the Declaration of Helsinki. Written informed consent was obtained from all patients before study entry.

### Patients and inclusion criteria

Inclusion criteria (all must be fulfilled) were: Patients with clinically non-granulomatous anterior uveitis with an AC cell grade >2+ according to SUN guidelines or with infectious endophthalmitis^1^. At the time of sampling patients received no topical anti-inflammatory medication. Standard laboratory parameters tested, and ophthalmic examinations are described in supplementary methods.

Patients included in the study were classified into the following groups (Suppl. Table. 1): 1) Patients with HLA-B27-associated acute anterior uveitis (B27+AAU), and with typical clinical signs of acute anterior uveitis. 2) Patients with HLA-B27 negative anterior uveitis (B27-AU), with typical clinical signs but without inflammatory / immune-mediated systemic disease associated with uveitis. 3) Patients with infectious endophthalmitis. Patients with ages, gender, and systemic medical therapies typical for these three entities were chosen.

### Laboratory parameters

The following standard laboratory parameters were tested in all patients: differential blood count, liver and kidney function tests, C-reactive protein, angiotensin-converting enzyme, soluble interleukin 2-receptor, serological testing for treponema pallidum, and the patient was excluded from the study if any of those were remarkable. Patients were analyzed for their HLA-B27 status using established PCR (licensed lab standard).

In addition, patients underwent chest x-ray and consultation with a specialist for internal medicine or rheumatology to identify any associated systemic immune-mediated disease. Patients were classified as HLA-B27-associated uveitis (eventually with associated systemic disease) if none of the tests except HLA-B27 positivity produced any further findings indicating non-related associated systemic disease. Patients with a clinical appearance of infectious (e.g., HSV-or VZV-induced) uveitis or uveitis syndromes (e.g., Fuchs uveitis syndrome) were not included in the study.

### Ophthalmic examinations

A standardized ophthalmic database was applied for the analysis, that included the following parameters: clinical ophthalmic observations on uveitis in the involved eyes were documented according to the SUN criteria^1^. Briefly, best-corrected visual acuity testing (in LogMAR), slit-lamp examination, Goldmann tonometry, and funduscopy were performed by two independent observers. Any uveitis-related intraocular complications were recorded (Suppl. Table. 1).

### AqH fine needle aspirates

AqH (100-150μl) was collected from each study subject using a 30G needle under local anesthesia, and immediately shipped at 4°C to the department of neurology at the University Clinic Muenster (Germany) for scRNA-seq and/or flow cytometry analysis. Cells were centrifuged once, counted and around 5,000 of the input cells were used for scRNA-seq, the remaining cells were used for flow cytometry (Method Details). For low input proteomics, 60 μl of the AqH were centrifuged for 5 min at 12,000g, and stored at −80° until analysis.

### Single Cell RNA-Sequencing and Analysis

Single cell suspensions were loaded onto the Chromium Single Cell Controller using the Chromium Single Cell 3’ Library & Gel Bead Kit v2 or v3 chemistry (both from 10X Genomics). Sample processing and library preparation was performed according to manufacturer’s instructions using AMPure XP beads (Beckman Coulter). Sequencing was either carried out on a local Illumina Nextseq 500 using the High-Out 75 cycle kit with a 26-8-0-57 read setup or commercially (Microanaly, China) on a NovaSeq 6000 using the 300 cycle kit with paired end 150 read setup. Samples library kits version and sequencing information are shown in Suppl. Tab. 2.

### Preprocessing of sequencing data

Processing of sequencing data was performed with the cellranger pipeline v3.0.2 (10x Genomics) according to the manufacturer’s instructions. Raw bcl files were de-multiplexed using the cellranger *mkfastq* pipeline. Subsequent reads alignments and transcript counting was done individually for each sample using the cellranger count pipeline with standard parameters. The cellranger *aggr* pipeline was employed to generate a single cell-barcode matrix containing all the samples without normalization. The normalization of each library was subsequently performed in Seurat (see below).

### Quality control, normalization, clustering, alignment and visualization of scRNA-seq datas

Subsequent analysis steps were carried out using Seurat v3.1.5^79^ using R v4.0.2 as recommended by the Seurat tutorials. Briefly, cells were filtered to exclude cell doublets and low-quality cells with few genes (<200), and high genes (>900-3500) or high mitochondrial percentages (5% – 7%) in each patient individually. After quality control, the total cell number used for the analysis was 12,305 (Suppl. Tab. 2). To account for technical variation, data were normalized using regularized negative binomial regression^80^ taking into account mitochondrial percentage and cycle score. Dimensionality reduction was done by Principal Component analysis. The number of principal components used for further analysis was determined using an elbow plot. Cells were clustered using the “FindNeighbors” (based on KNN graphs) and “FindCluster” (Louvain algorithm) function in Seurat. To account for batch effects, different samples were aligned using Harmony^81^. The UMAP was then used to visualize cells in a two-dimensional space. Clusters were annotated based on known marker genes.

### Identifying differentially expressed genes

The “FindMarker” function in Seurat, which used the Wilcoxon rank sum test, was applied to normalized and aligned data. The threshold of the adjusted p value was set to 0.05. Volcano plots were created with the R package *EnhancedVolcano*. Differentially expressed genes identified by Seurat were used as input. The threshold for the average log fold change was set at 0.5 and for p values at 0.001.

### Average expression of GWAS risk gene

Summary statistics were downloaded from the NHGRI-EBI GWAS Catalog^82^ for the studies GCST007362/GCST007361^83^, GCST001345^84^, GCST000563^85^, GCST007361^83^, GCST007844^86^, GCST001149^87^, GCST005529^88^, GCST008910^89^, GCST003097^90^, and GCST010481^91^ downloaded on 04/09/2020.

### Identifying cellular interactions

Cellular interactions were analyzed using CellPhoneDB^43^. Normalized and aligned scRNA-seq data with the clusters identified by Seurat separated by diagnosis were used for analysis. Clusters with less than 10 cells were excluded. Statistical iterations were set at 1000 and genes expressed by less than 10% of cells in a cluster were removed. Resulting interactions are based on the CellPhoneDB repository. Heatmaps were produced by using the integrated heatmap function, and then calculating the difference of the count of significant interactions in condition1 and condition2. Dotplots were created with the integrated dotplot function.

### Flow cytometry of AqH -derived cells

Flow cytometry analysis was performed on the maximum of recovered cells from the AqH samples, (≤10^6^ cells). Cells were first blocked with FcR anti-human blocking reagent (Miltenyi) Afterwards, cells were stained for 30 min at 4°C in the dark with a combination of anti-human antibodies: CD3 (perCp-Cy5.5, clone OKT3), CD4 (BV510, clone OKT4), CD8 (APC, clone SK1), CD11b (FITC clone M1/70), CD11c (Pacific Blue, clone 3.9), CD56 (Pe-Cy7, clone N901) _ all from Biolegend _ and HLA-DR (ECD, clone Immu-357) from Beckman Coulter. Samples were measured on a Gallios (10 colors, 3 lasers, Beckman Coulter) flow cytometer using FACS Kaluza software v2.1.1 (Beckman Coulter). Data were analyzed with FlowJo v10.6.1 (BD Biosciences). The gating strategy is illustrated in Suppl. Fig. 3.

### Quantification of cytokines in AqH via low input proteomics

Cytokines in AqH samples were quantified using a ProcartaPlex Human Cytokine-Panel 1B (Thermo Fisher Scientific, Waltham Massachusetts, USA) that quantifies: granulocyte macrophage-colony stimulating factor (GM-CSF), interferon (IFN)-α, IFN-γ, interleukin (IL)-1α, IL-1β, IL-1RA, IL-2, IL-4, IL-5, IL-6, IL-7, IL-9, IL-10, IL-12p70, IL-13, IL-15, IL-17A, IL-18, IL-21, IL-22, IL-23, IL-27, IL-31, tumor necrosis factor (TNF)-α, and TNF-β), according to the manufacturer’s instructions. Standards and samples were measured in duplicates using Bio-Plex MAGPIX Multiplex Reader (BioRad, Hercules, California, USA) and cytokines were quantified in [pg/μl] using ProcartaPlex Analyst 1.0 software (Thermo Fisher Scientific).

## Abbreviations

B27-AU: HLA-B27 negative anterior uveitis;
B27+AAU: HLA-B27 positive acute anterior uveitis;
AqH: aqueous humor,
AC: anterior chamber,
EIU: endotoxin-induced uveitis.

## Acknowledgements

G. M.z.H. was supported by grants from the Deutsche Forschungsgemeinschaft (DFG, ME4050/4-1, ME4050/7-1), from the Innovative Medical Research (IMF) program of the University Münster, from the Multiple Sclerosis Innovation (Merck), and from the Ministerium für Innovation, Wissenschaft und Forschung (MIWF) des Landes Nordrhein-Westfalen. A. H. was supported in part by the German Society of Ophthalmology (DOG). We thank Dr. Carsten Heinz for patient recruitment and Dr. Susanne Wasmuth for technical assistance.

## Author Contributions

M.K., D.S., X.L., T.L., D.B., K.L. performed experiments, M.H., K.W. performed computational analyses, G.M.z.H., A.H., and M.K. conceived the study. G.M.z.H., and A.H. supervised the study. H. W. co-supervised the study. G.M.z.H., M.K., M.H., and A.H. wrote the manuscript. All authors critically revised the manuscript and agreed with its contents.

## Data and Code Availability

Raw sequencing data are available in the Gene Expression Omnibus (GEO) repository (accession code will be made public upon manuscript publication). We followed the official tutorial of the packages listed and did not generate any custom code. An interactive version of the scRNA-seq data was created using cerebroApp^92^ and is available at: http://uveitis.mheming.de.

## Supplementary Figure Legends

**Suppl. Fig. 1.**
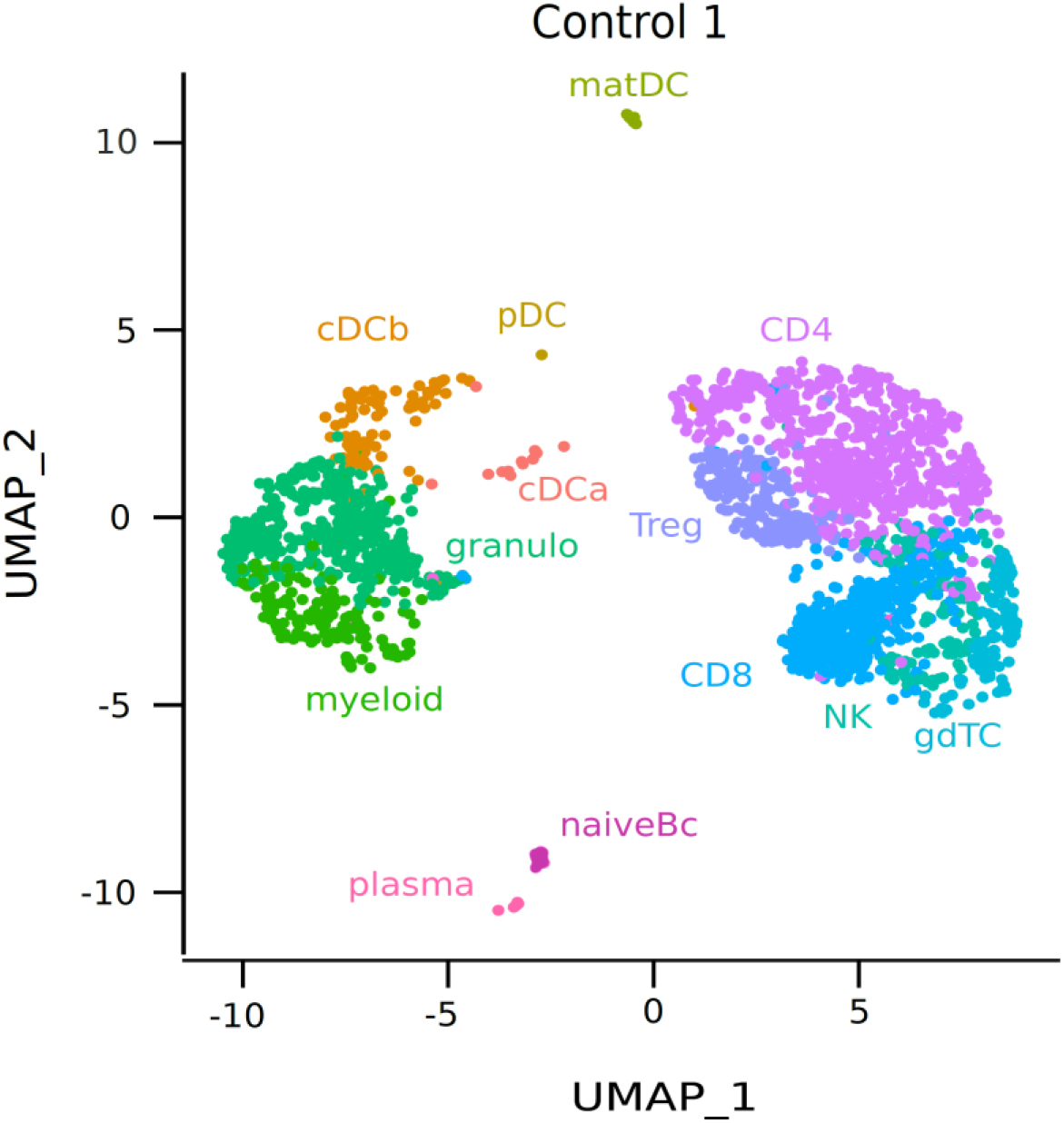
Individualized scRNA-seq results of anterior chamber-derived leukocytes. UMAP projection as in Fig. 2A of only the endophthalmitis control patient. The sc transcriptomes were manually annotated to cell types based on marker gene expression, and distinguished in 13 cell clusters (color-coded; each dot represents one cell).

**Suppl. Fig. 2.**
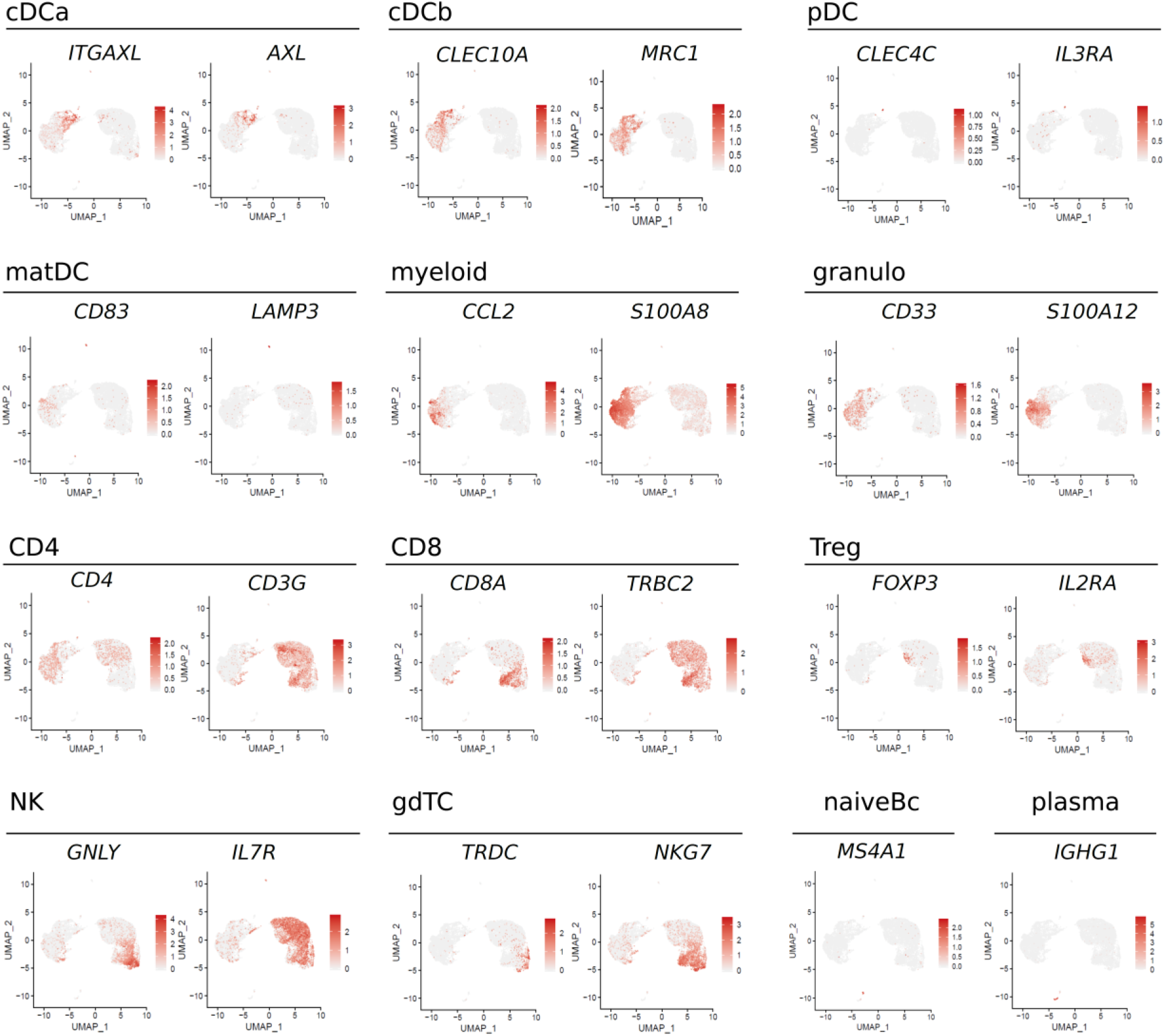
Feature plots of lineage marker genes. Expression of lineage marker *ITGAX, AXL, CLEC10A, MRC1, CLEC4C, IL3RA, CD83, LAMP3, CCL2, S100A8, CD33, S100A12, CD4, CD3G, CD8A, TRBC2, FOXP3, IL2RA, GNLY, IL7R, TRDC, NKG7, MS4A1, IGHG1* are shown as feature plots. Each dot represents one cell.

**Suppl. Fig. 3.**
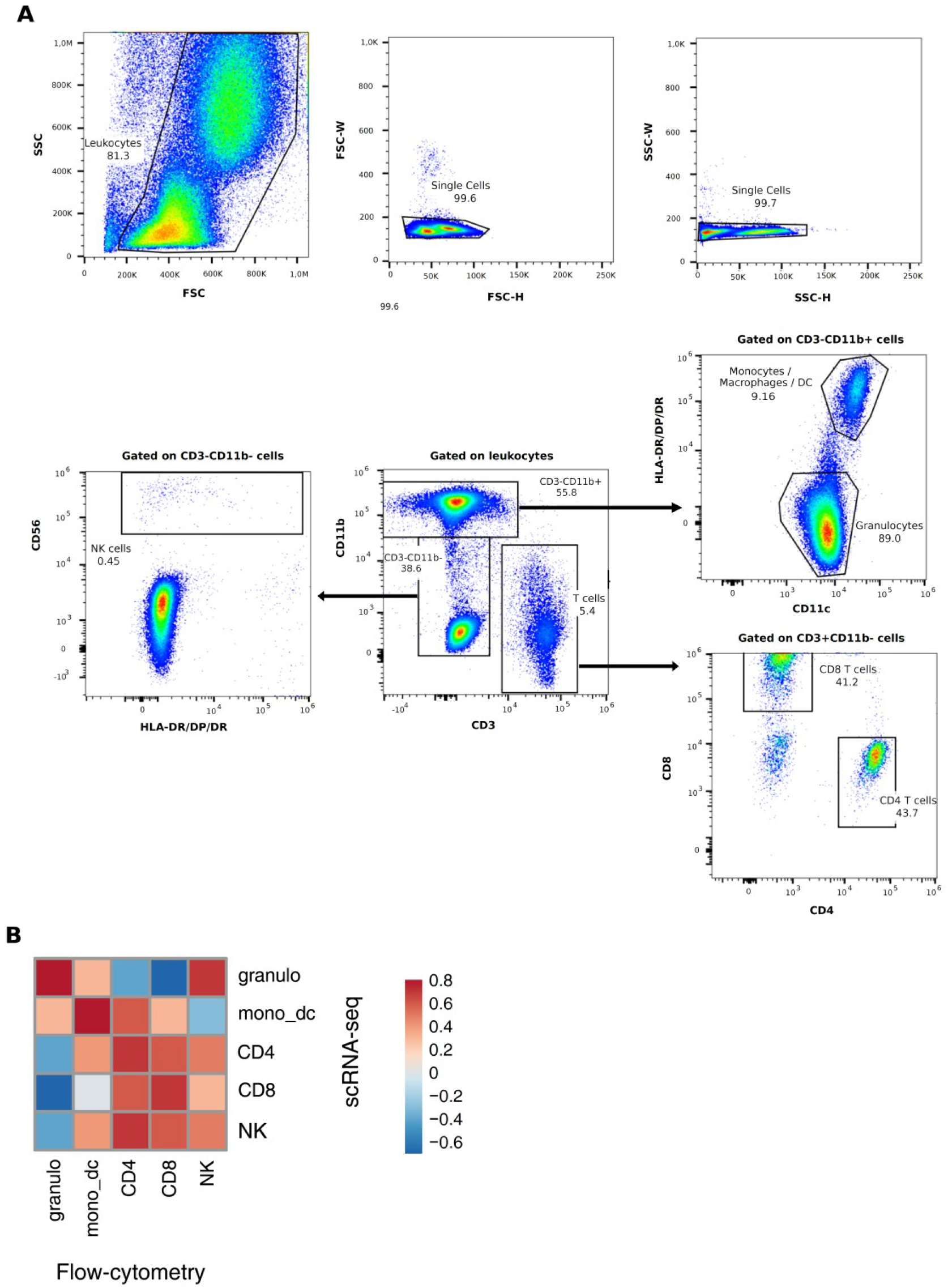
Flow cytometric analysis of aqueous humor samples. (A) Leukocytes in SSC/FSC scatter were identified, doublets were excluded, and leukocytes were gated on CD3-CD11b+ myeloid cells, CD3+CD11b-T-cells and CD3-CD11b-cells. Myeloid cells were further classified in CD11c+HLA-DR-granulocytes and CD11c+HLA-DR+ monocytes / macrophages / DC. T-cells were further subdivided in CD4+ and CD8+ expressing cells. CD3-CD11b-cells were analyzed according to their frequency of CD56+ NK-cells. Representative analysis is shown. (B) Correlation plot (Spearman’s correlation coefficient) between flow proportions (columns) and scRNA-seq (rows) proportions are shown as a heatmap with high correlation coefficients shown in red.

**Suppl. Fig. 4.**
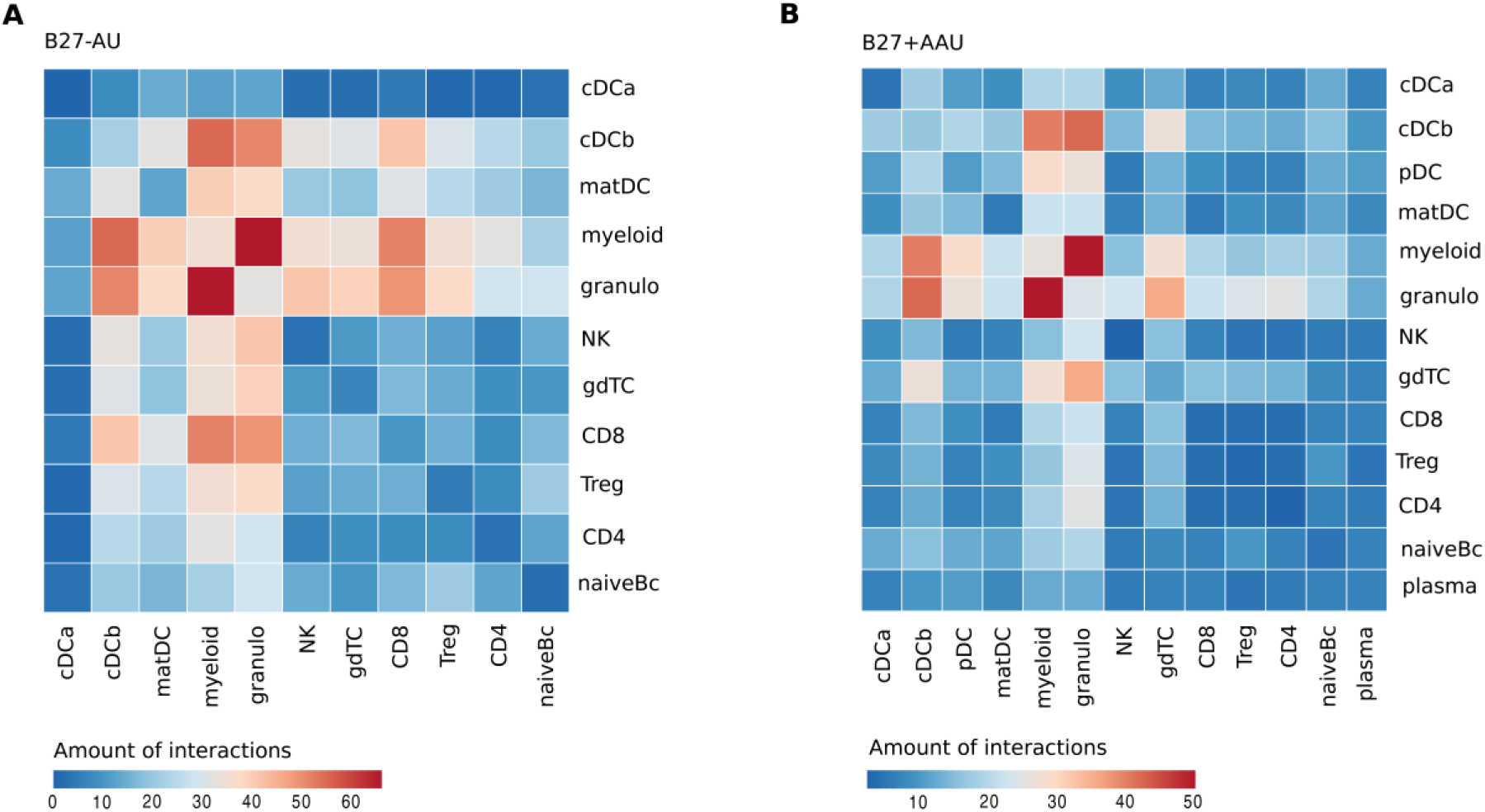
Detailed results of the interactome prediction analysis. Predicted receptor-ligand interactions between cell clusters in (A) B27-AU and (B) B27+AAU were obtained with CellPhoneDBv2.0. The legend represents the amount of interactions.

**Suppl. Fig. 5.**
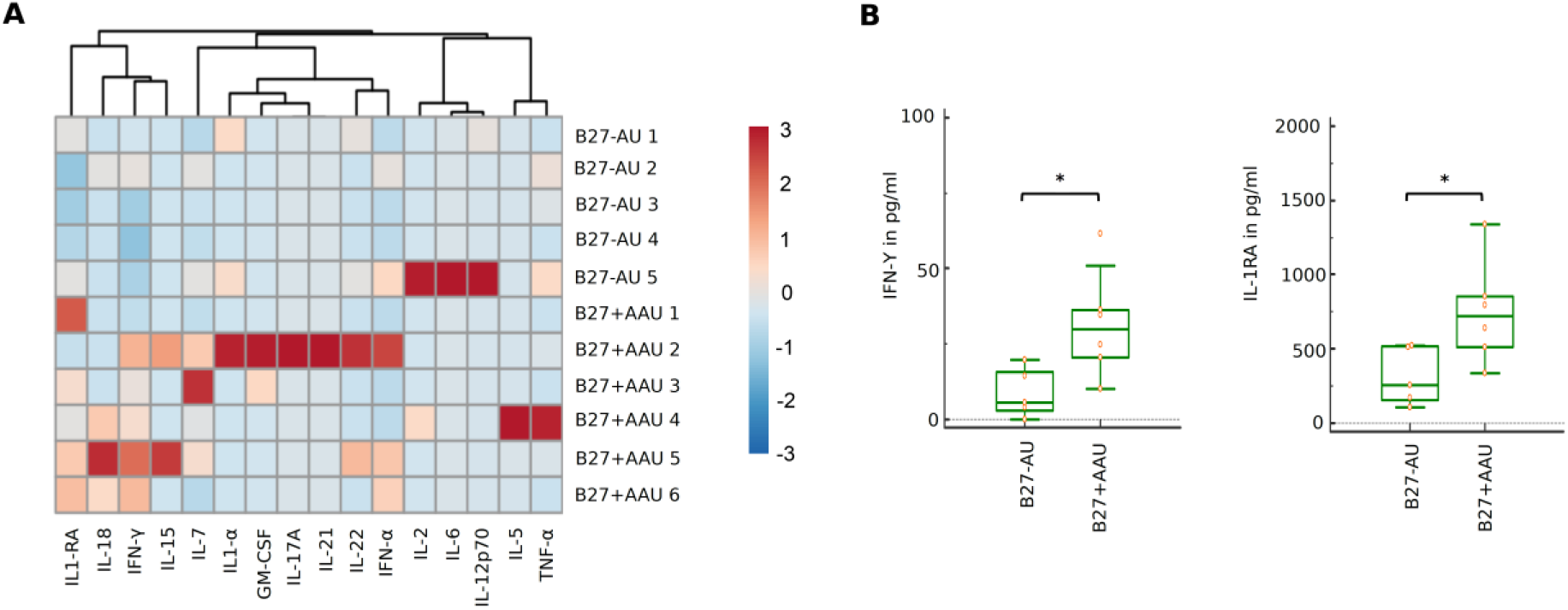
Proteomics analysis in serum. (A) Heatmap showing serum cytokine level of B27-AU (n=5) and B27+AAU (n=6) patients (see Suppl. Tab.9 showing single values). Data were scaled row wise. Columns are clustered hierarchically using euclidean distance measure and complete linkage. (B) Box plots of IFN-γ, and IL-1RA (pg/ml) in the sera of patients with B27-AU and B27+AAU. Dots represent individual data. Mann-Whitney U-test (* p<0.05).

## Supplementary Table Legends

Supplementary Table 1: Clinical data and laboratory analysis

Supplementary Table 2: Summary of technical information regarding libraries preparation and sequencing

Supplementary Table 3: List of top DE genes per cluster in Figure 2

Supplementary Table 4: Absolute and relative cluster size in Figure 2

Supplementary Table 5: List of DE genes per meta cluster in Figure 3

Supplementary Table 6: List of the DE genes per cluster in Figure 2

Supplementary Table 7: GWAS risk genes per cluster in Figure 2 and Figure 3G

Supplementary Table 8: AqH Cytokine level in Figure 4

Supplementary Table 9: Serum cytokine level in Supplementary Figure 5

